# Towards a biotechnological platform for the production of human pro-angiogenic growth factors in the green alga *Chlamydomonas reinhardtii*

**DOI:** 10.1101/831933

**Authors:** Montserrat Jarquín-Cordero, Myra Noemi Chávez, Carolina Centeno-Cerdas, Alexandra-Viola Bohne, Ursula Hopfner, Hans-Günther Machens, José Tomás Egaña, Jörg Nickelsen

## Abstract

The recent use of photosynthetic organisms such as *Chlamydomonas reinhardtii* in biomedical applications has demonstrated their potential for the treatment of acute and chronic tissue hypoxia. Moreover, transgenic microalgae have been suggested as an alternative *in situ* drug delivery system. In this study, we set out to identify the best available combination of strains and expression vectors to establish a robust platform for the expression of human pro-angiogenic growth factors, i.e. hVEGF-165, hPDGF-B, and hSDF-1, in biomedical settings. As a case study, combinations of two expression vectors (pOpt and pBC1) and two *C. reinhardtii* strains (UVM4 and UVM11) were compared with respect to hVEGF-165 transgene expression by determination of steady-state levels of transgenic transcripts as well as immunological detection of recombinant proteins produced and secreted by the generated strains. The results revealed the combination of the UVM11-strain with the pBC1-vector to be the most efficient one for high level hVEGF-165 production. To assess the robustness of this finding, the selected combination was used to create hPDGF-B and hSDF-1 transgenic strains for optimized recombinant protein expression. Furthermore, biological activity and functionality of algal-produced recombinant pro-angiogenic growth factors were assessed by receptor phosphorylation and *in-vitro* angiogenesis assays. The results obtained revealed a potentiating effect in the combinatorial application of transgenic strains expressing either of the three growth factors on endothelial cell tube formation ability, and thus support the idea of using transgenic algae expressing pro-angiogenic growth factors in wound healing approaches.

## Introduction

Ever since *C. reinhardtii* was discovered as an economically attractive and sustainable green microbial cell factory, the list of its applications has grown as its many advantages have been recognized (Rasala et al. 2014; Rasala and Mayfield 2015; Dyo and Purton 2018). As a unicellular photosynthetic organism, *C. reinhardtii* is in many respects easy to grow in culture. Growth requirements are comparatively simple, well defined and cost-effective, which makes scaling up straightforward. Also, since *C. reinhardtii* is a well-studied model organism, its three genomes have been fully sequenced and an advanced molecular toolkit has been developed that facilitates the construction of transgenic algae (Rasala and Mayfield 2011). Taking advantage of these methods, we explored the use of genetically modified *C. reinhardtii* strains that are capable of the simultaneous production and local delivery of both oxygen and bioactive molecules that promote wound healing in biomedical settings. In the context of this novel approach, named HULK (from ‘*Hyperoxie Unter Licht Konditionierung’*, the German term for ‘light-induced hyperoxia’; Hopfner et al. 2014; Schenck et al. 2015), three transgenic *C. reinhardtii* strains that express and secrete different functional human pro-angiogenic growth factors were generated (Chávez et al. 2016; Centeno-Cerdas et al. 2018). Their basic ability to produce oxygen through the local induction of photosynthesis, as well as sustainably release human vascular endothelial growth factor A (hVEGF-165), platelet-derived growth factor B (hPDGF-B) and stromal cell-derived factor 1 (hSDF-1) has been demonstrated by combining these transgenic algae with biomaterials, such as dermal scaffolds and non-absorbable sutures, both *in vitro* and *in vivo* (Chávez et al. 2016; Centeno-Cerdas et al. 2018). However, the significant differences in expression levels of each recombinant protein revealed a need for further optimization of the strategy.

On the one hand, recombinant protein expression based on nuclear transgenes in *C. reinhardtii* is particularly convenient when compared to other systems. Nuclear transformation of this alga is readily achieved by a simple protocol that inserts the desired DNA into the host’s nuclear genome. Furthermore, as a eukaryotic organism, the alga itself possesses the machinery required to mediate important post-translational modifications, such as the formation of disulfide bridges, and nitrogen (N-) and oxygen (O-) linked glycosylation, which are relevant to the functionality of the desired product (Mathieu-Rivet et al. 2017). Moreover, the discovery of several subcellular targeting signals directly enables recombinant protein secretion into the medium, thus avoiding the need for expensive or tedious isolation protocols (Ramos-Martinez et al. 2017; Molino et al. 2018; Baier et al. 2018). However, one major drawback that limits the use of *C. reinhardtii* as an expression platform for recombinant growth factors is the low level of product obtainable from nuclear transgenes – a problem that has persisted despite more than 20 years of research (Fuhrmann et al. 1999). Several key studies have addressed the potential impact of the genetic background of *C. reinhardtii* strains, as well as the influence of regulatory sequences such as promoters, introns and terminator sequences, on the efficiency of recombinant protein expression. Of these studies, three are of particular importance for the development of the project presented here. The respective reported strains and expression vectors were chosen because of their promising potential to become elements of an efficient and versatile *C. reinhardtii* expression platform.

The first of these papers described a vector system based on the nuclear *C. reinhardtii PsaD* gene and its regulatory elements, which was shown to support high-level expression of both endogenous and exogenous genes (Fischer and Rochaix 2001). Moreover, this vector was described to allow the efficient generation of transgenic strains as well as the subcellular targeting of recombinant proteins. In the second study, Neupert and co-workers implemented a strategy designed to select *C. reinhardtii* mutants that were able to express transgenes efficiently and accumulated recombinant proteins to high levels (Neupert et al. 2009). By assuming that poor transgene expression was related to a genetically based suppression mechanism, these authors isolated UV light-induced mutant clones that were deficient for it. They generated two cell-wall-deficient *C. reinhardtii* strains, UVM4 and UVM11, each of which displayed high transformability, expressed nuclear transgenes at high levels and produced large amounts of recombinant proteins. By using a vector containing a synthetic GFP gene whose codon usage was optimized for *C. reinhardtii* (Fuhrmann et al. 1999) and which was expressed under the control of the *PsaD* promoter and terminator elements described above (pJR38/pBC1-CrGFP), high recombinant protein yields of up to 0.2 % of the total soluble protein were achieved. Finally, in the third study, Lauersen and co-workers described the development of a vector system that secreted recombinant proteins at levels of up to 10 mg·ml^−1^ (Lauersen et al. 2013). It was later optimized by the same group to become a modular vector (pOpt) that allows rapid cloning of genes of interest for nuclear expression in *C. reinhardtii* (Lauersen et al. 2015). The expression cassette in this construct uses the Heat Shock 70A (P_*HSP70A*_)-Rubisco small subunit 2 (P_*RBCS2*_) fusion promoter, together with the Rubisco small subunit intron 1 (*RBCS2-i1*) and the 3’ untranslated region of *RBCS2* as regulatory elements (Schroda et al. 2000).

In this work, these three promising approaches were implemented in a search for the optimal combination of algal strain and expression vector that would lead to significantly enhanced yields of the recombinant human growth factors we had previously expressed in *C. reinhardtii*. The results obtained show how promoters, signal peptides and purification tags – even those that have been reported to work very well with other reporter genes – can greatly affect recombinant protein yields. A significant advantage of using the UVM11 strain was documented, as the newly created transgenic strains outperformed the previous UVM4-derived strains with respect to expression yields and functionality of the recombinant proteins. Finally, based on an *in vitro* angiogenesis assay, the augmented biological effect of a combination of transgenic strains expressing different human growth factors was demonstrated.

## Materials and Methods

### *C. reinhardtii* cell culture

The cell-wall-deficient (*cw15) C. reinhardtii* strains UVM4 and UVM11 (Neupert et al. 2009) and derived transgenic strains were grown mixotrophically at 23°C on either solid Tris Acetate Phosphate (TAP) medium or in liquid TAP medium supplemented with 1 % (w/v) sorbitol (TAPS) (Harris 2008) under standard culture conditions (23°C, constant illumination at 30 μE·m^−2^·s^−1^ and agitation at 120 rpm).

### pBC1 vector derivatives

The pBC1-CrGFP (pJR38, Neupert et al. 2009)-derived vectors pBC1_VEGF-165, pBC1_PDGF-B and pBC1_SDF-1, which were described previously (Chávez et al. 2016; Centeno-Cerdas et al. 2018) and have been used to create *C. reinhardtii* UVM4 transgenic strains secreting the human growth factors hVEGF-165 (pBC1_V-4), hPDGF-B (pBC1_P-4) and hSDF-1 (pBC1_S-4), respectively, were transformed into the UVM11 strain (pBC1_V-11, pBC1_P-11, pBC1_S-11). Briefly, the synthetic human genes with sequences adapted to the codon usage of *C. reinhardtii* (GenBank Accession No.: MN496135, MN496136, MN496137) were cloned into the pBC1-CrGFP vector backbone via its *Nde*I and *EcoR*I restriction sites, thereby replacing the CrGFP cassette between the endogenous *PsaD* 5’ and 3’ UTRs (Fig. 1). For secretion of the recombinant proteins, the sequence encoding the 21-amino-acid leader peptide of the *C. reinhardtii* extracellular enzyme arylsulfatase (ARS2, Cre16.g671350, Phytozome, release C. reinhardtii v5.5) was inserted upstream of the protein-coding sequence. The vectors also carried the *APH*VIII resistance gene, providing for selection on paromomycin.

**Figure 1:**
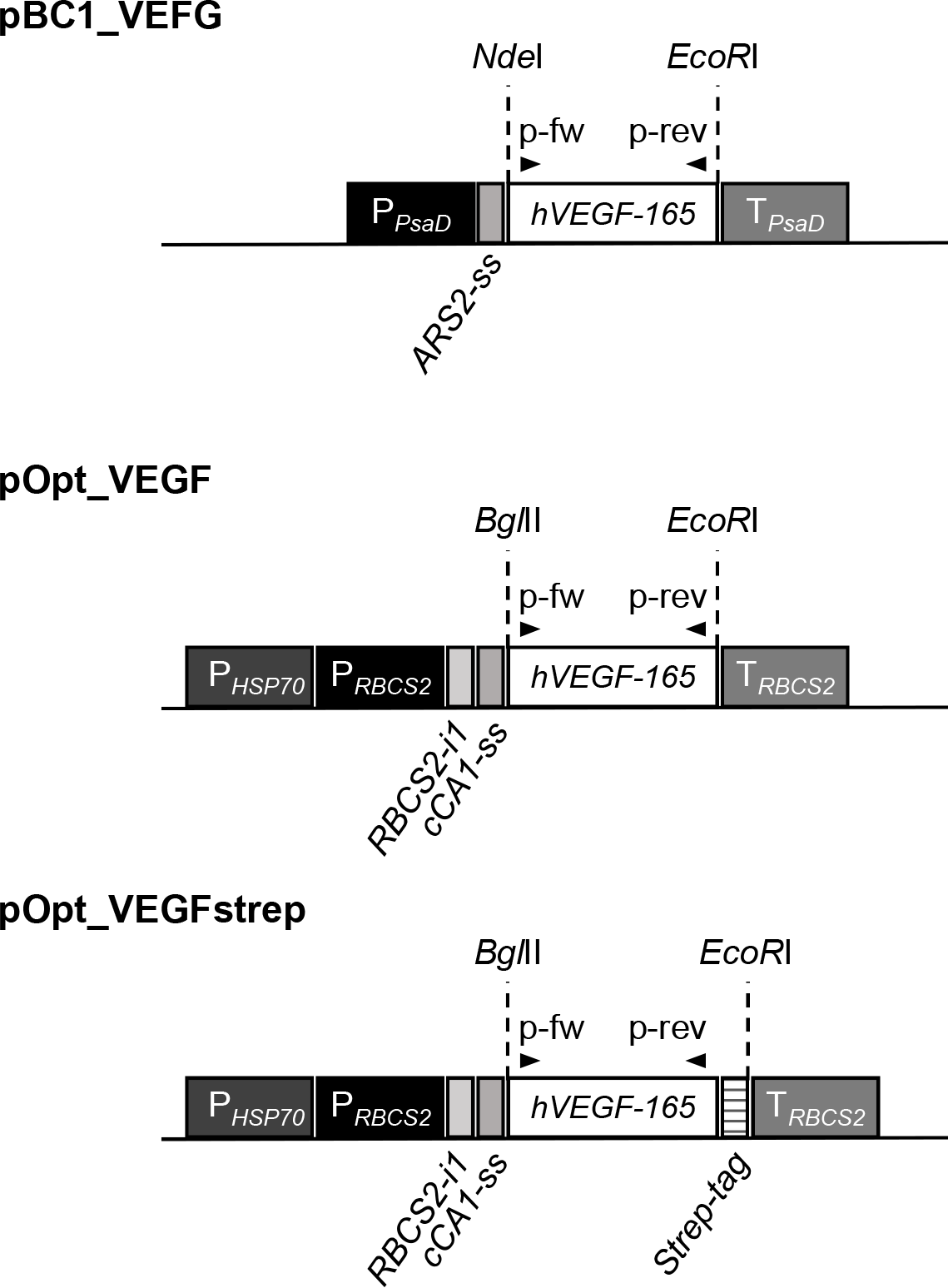
Schematic representation of vectors created to generate hVEGF-165 expressing transgenic *C. reinhardtii* strains. The hVEGF-165 transgene in the pBC1_VEGF-165 construct was placed under the control of the *PsaD* promoter (P_*PsaD*_) and terminator (T_*PsaD*_) regions. The regulatory elements in the pOpt_V expression cassette comprised the hybrid heat shock protein 70A-Rubisco small subunit 2 promoter P_*HSP70*_-P_*RBCS2*_, the *RBCS2* intron (*RBCS2-i*) and terminator sequences (T_*RBCS2*_). Protein targeting into the culture medium was achieved by adding secretion signals derived from *C. reinhardtii* arylsulfatase 2 (ARS2) or carbonic anhydrase 1 (cCA1) upstream of the hVEGF-165 coding sequence, respectively. The positions of primers used for PCR analysis to verify the integration of the hVEGF-165 gene into the *C. reinhardtii* nuclear genome are indicated (p-fwd: forward primer, p-rev: reverse primer).

### pOpt vector derivatives

The coding sequence for synthesis of hVEGF-165 (GenBank Accession No. NP_001165097) was amplified by PCR from the construct pBC1_VEGF-165, using synthetic primers to add a stop codon and restriction sites for *Bgl*II and *EcoR*I to enable their insertion into the basis vector pOpt_mVenus_Paro (Lauersen et al. 2015). The PCR product was then inserted into the pOpt expression cassette between the *HSP70A-RBCS2-i1* promoter and the *RBCS2* 3’ UTR, and downstream of the sequence encoding the 23-amino-acid leader peptide of *C. reinhardtii* periplasmic enzyme carbonic anhydrase 1 (cCA1, Cre04.g223100, Phytozome, release *C. reinhardtii* v5.5) that enables the proteins to be secreted into the culture medium. The resulting construct was named pOpt_VEGF-165. To create the construct pOpt_VEGF-165-strep (pOpt_VEGF-165-s) the coding sequences were amplified using PCR primers that added the same restriction sites but obliterated the stop codon sequence, thus allowing for translation of the sequence encoding the C-terminal Strep tag (amino acid sequence: WSHPQFEK). The sequences of the primers used for cloning experiments were VEGF-*Bgl*II-fw 5’-AGATCTGCCCCCATGGCCGAGGGC-3’; VEGF-*EcoR*I-rev 5’-GAATTCTTAGCGGCGGGGCTTGTCG-3’; VEGF-*EcoR*I-rev2: 5’-GAATTCTAAGCGGCGGGGCTTGTCG-3’. All constructs were verified by sequencing prior to *C. reinhardtii* transformation.

### Nuclear transformation of *C. reinhardtii*

*C. reinhardtii* UVM4 or UVM11 cultures were grown to mid-log phase under standard conditions. Then, 10^7^ cells were suspended in a volume of 1 ml and vortexed with glass beads (diam. 0.5 mm) for 20 s in the presence of 5 μg of plasmid DNA. Following incubation overnight under low light levels, the cells were grown under standard levels of illumination on TAP-Agar plates containing paromomycin (10 mg·ml^−1^) to select for transgenic clones.

### PCR assays

Genomic DNA was extracted from 50 ml of *C. reinhardtii* cells using phenol/chloroform/isoamyl alcohol (25:24:1) (Carl Roth GmbH, Mannheim, Germany) and chloroform/isoamyl alcohol (24:1). DNA was then precipitated with ice-cold isopropanol and washed twice with 70% ethanol. Pellets were air-dried for 5 min and resuspended in 50 μL H_2_O. Integration of the recombinant genes was confirmed by polymerase chain reaction (PCR) using the following transgene-specific primer pairs (Metabion GmbH, Planegg, Germany): 5’-GAAGTTCATGGACGTGTACC-3’ and 5’-TTGTTGTGCTGCAGGAAG-3’ for hVEGF-165 (258-bp product); 5’-AACGCCAACTTCCTGGTG-3’and 5’-GTGGCCTTCTTGAAGATGGG-3’ for hPDGF-B (164-bp product); 5’-CGTGAAGCACCTGAAGATCC-3’ and 5’-CTTCAGCTTGGGGTCGATG-3’ for hSDF-1 (103-bp product).

### Southern blot

Genomic DNA was extracted from 150-ml cultures of each strain using CTAB buffer (2 % cetyltrimethylammonium bromide, 100 mM Tris-HCl, pH 8,) and phenol/chloroform/isoamyl alcohol (25:24:1) (Carl Roth GmbH, Mannheim, Germany). Aliquots (30 μg) of DNA were then digested with *Xho*I or *BamH*I at 37 °C for 48 h. Samples were analyzed as described before (Chávez et al. 2016), using gene-specific digoxigenin-nucleotide-labelled DNA probes (Digoxigenin-11-dUTP alkali-labile, Roche, Basel, Switzerland) obtained using the same primer pairs and conditions as for the PCR. Signals were detected using an alkaline-phosphatase-conjugated anti-DIG antibody (Roche, Basel, Switzerland) and CDP* (Roche) as reaction substrate.

### Northern blot

To analyze transcript accumulation in the different strains expressing hVEGF-165, 50-ml samples of *C. reinhardtii* cells were harvested at mid-log phase (~3 ×10^6^ cells·ml^−1^) by centrifugation (4000 g, 15 min, 4°C) and total cellular RNA was extracted using Tri-Reagent^®^ (Sigma Aldrich, Darmstadt, Germany). Aliquots (40 μg) of total RNA were electrophoretically fractionated on denaturing agarose gels, blotted onto positively charged nylon membranes (Roti^®^-Nylon plus, pore size 0.45 μm; Carl Roth GmbH, Karlsruhe, Germany), hybridized to gene-specific digoxigenin-nucleotide labelled DNA probes and visualized as described above.

### Protein isolation and Western blot analyses of the recombinant protein

To quantify the level of expression of the hVEGF-165 protein, *C. reinhardtii* cells were inoculated in triplicate into 100 ml of TAPS and incubated under standard conditions for approximately 4 to 5 days or until they reached a density of 10^7^ cells·ml^−1^. Supernatants were collected and passed through a 0.22-μm filter (ClearLine^®^, Kisker Biotech GmbH & Co. KG, Steinfurt, Germany), and then centrifuged (3000 g, 40 min) through Amicon^®^Ultra-15 30K filter units (Merck Millipore Ltd., Carrigtwohill, Ireland) to concentrate the recombinant protein to a final volume of 250 μl. Protein amounts were determined with the Bradford assay (Roti^®^-Quant, Carl Roth GmbH) according to the manufacturer’s instructions. Then, 15-μg aliquots of protein were denatured at 95 °C for 5 min in the presence of reducing loading buffer Roti^®^Load-1 (Roth GmbH), fractionated by SDS-PAGE (12 % acrylamide), and transferred to nitrocellulose membranes (0.45 μm, AppliChem GmbH, Darmstadt, Germany). Commercially available recombinant hVEGF-165 was loaded in parallel as a positive control (Peprotech, Rocky Hill, NJ, USA). Protein detection was performed using a monoclonal rabbit anti-VEGF primary antibody (1:1000 dilution, ab52917, Abcam plc, Cambridge, UK) and a goat anti-rabbit HRP secondary antibody (1:5000, Dianova GmbH, Hamburg, Germany), using the SuperSignal West Pico detection system (Pierce-Thermo Fisher Scientific Inc., Rockford, IL, USA).

### Enzyme-linked immunosorbent assay (ELISA) for the detection of recombinant growth factors

*C. reinhardtii* cells were inoculated in triplicate into 25 ml TAPS-liquid cultures and incubated under standard conditions for approximately 4 to 5 days or until they reached a cell density of 10^7^ cells·ml^−1^. Culture supernatants were collected by centrifugation (5 min, 10,000 g) and then stored at −80 °C before analysis. The pelleted cells were then resuspended to a density of 10^7^ cells in 200 μl of lysis buffer (100 mM Tris-HCl, 10 mM EDTA, 0.5 % Triton-X-100, 25 mg·ml^−1^ pepstatin, 25 mg·ml^−1^ leupeptin, 25 mg·ml^−1^ aprotinin) and disrupted by vortexing with glass beads (diam.: 0.5 mm). The protein concentration in the supernatant was determined by the Bradford assay (Roti^®^Quant, Carl Roth GmbH), and with the Pierce™ BCA Protein assay (Thermo Scientific) in the protein lysates, both according to the manufacturer’s instructions. For recombinant protein quantification in the medium samples (supernatant) and lysates (cells), the human VEGF DuoSet ELISA kit, human PDGF-BB DuoSet ELISA kit and human CXCL12/SDF-1 DuoSet ELISA kit (R&D Systems, Minneapolis, MN, USA) were used according to the manufacturer's instructions. The secretion and retention ratios were calculated from the normalized growth factor concentrations determined in the supernatant and the cell lysate, respectively (total concentration of growth factor (100%) = growth factor concentration in cell culture supernatant pro μg total protein + growth factor concentration in cell lysate pro μg total protein).

### Recovery of recombinant proteins from the culture supernatants

Triplicate samples of *C. reinhardtii* strains were cultured under standard conditions in 150 ml TAPS-liquid cultures until they reached a density of 10^7^ cells·ml^−1^. In order to obtain protein concentrates with comparable amounts of the different recombinant growth factors, based on previous experience, different volumes of culture supernatant were filtered and concentrated (30 ml pBC1_V-4 and pBC1_V-11, 60 ml pOpt_V-4 and pOpt_V-11, 150 ml pOpt_Vs-4 and pOpt_Vs-11, 60 ml pBC1_P-4 and pBC1_P-11 and 150 ml pBC1_S-4 and pBC1_S-11). Filtration units capable of retaining peptides with molecular weights above 30 kD, Amicon^®^Ultra15 30K (Merck Millipore Ltd., Carrigtwohill, Ireland) were used to recover hVEGF-165 and hPDGF-B, while the Amicon^®^Ultra-15 10K (Merck Millipore Ltd., Carrigtwohill, Ireland) filter tubes were used to recover hSDF-1. Then, in a final filtration step, the diluent was changed to cell starvation medium [AIM-V serum-free medium with stable glutamine, streptomycin sulfate (50 μg ·ml^−1^) and gentamycin sulfate (10 μg·ml^−1^), Life Technologies, Grand Island, NY, USA) for HUVECs, or RPMI 1640 (with stable glutamine and 2.0 g·L^−1^ NaHCO_3_, Biochrom, Berlin, Germany) supplemented with 1 % fetal calf serum (heat-inactivated FCS, Biochrom) for hASC]. The total protein content of this conditioned medium was assessed by Bradford quantification (Roti^®^Quant, Carl Roth GmbH), whereas the amount of recombinant protein (hVEGF-165, hPDGF-B and hSDF-1) was quantified by ELISA.

### Cell culture of human cells

Human umbilical vein endothelial cells (HUVECs) were purchased (Promocell, Heidelberg, Germany) and maintained in supplemented Endothelial Cell Growth Medium 2 (Promocell) with 1 % penicillin/streptomycin (Biochrom) under standard cell culture conditions (37 °C, 5 % CO_2_). For all experimental settings, cells from passages 2-5 were used. Human adipose-derived stem cells (hASCs) were purchased (PT-5006, Lonza, Basel, Switzerland), and maintained in StemMACS™ MSC Expansion Media (Miltenyi Biotec GmbH, Bergisch Gladbach, Germany) supplemented with 1 % antibiotic/antimycotic (100× ab/am; Capricorn Scientific, Ebsdorfergrund, Germany). For all experimental settings, cells from passages 2 to 5 were used.

### Western blot receptor phosphorylation assay

To evaluate the biological activity of hVEGF-165, a receptor phosphorylation assay was performed. For this purpose, 1·10^5^ HUVECs per well were cultured for 24 h in 12-well plates and then starved for 16 h before activation by cultivation in AIM-V serum-free medium. Cells were then stimulated for 5 min, with either 50 ng·ml^−1^ recombinant hVEGF-165 (Peprotech, Rocky Hill, NJ, USA) or concentrated protein supernatants obtained from cultures of the genetically modified or recipient (UVM4/UVM11) strains containing either the same amount of total protein or the same amount of recombinant hVEGF-165. Cells were then snap-frozen by submerging the plate in liquid nitrogen and lysed in RIPA buffer (10 mM Tris-HCl, 100 mM NaCl, 0.5 % NP-40, 0.5 % deoxycholic acid, 10 mM EDTA) containing phosphatase inhibitors (Phosphatase Inhibitor Mini Tablets; Pierce-Thermo Fisher Scientific) and proteinase inhibitors [PIC, BD Pharmingen, Franklin Lakes, NJ, USA; Pefabloc SC-Protease Inhibitor, Carl-Roth, Karlsruhe, Germany; cOmplete™, Roche, Basel, Switzerland; and PMSF, Sigma Aldrich, St. Louis, MO, USA) in the concentrations recommended by the relevant manufacturer. Cells were scraped from the well and lysates were homogenized by pipetting up and down and stored at −80 °C for further analysis. For Western blot analysis, equal protein amounts were loaded onto 7.5% polyacrylamide gels (Mini-PROTEAN^®^TGX Stain-Free™ Precast Gels, Bio-Rad Laboratories, Hercules, CA, USA) and fractionated by gel electrophoresis under reducing conditions and then blotted onto PVDF membranes (Immun-Blot^®^PVDF Membrane, Bio-Rad Laboratories). The primary monoclonal antibodies rabbit mAb anti-VEGFR-2 (55B11) (Cell Signaling Technology, Danvers, MA, USA) and rabbit mAb anti-phospho-VEGFR-2 (Tyr1175) (Cell Signaling), and the secondary goat-anti-rabbit (1:5000, Dianova GmbH) were used for detection of the phosphorylated and non-phosphorylated receptor epitopes, respectively, with overnight incubation periods. Clarity™ Western ECL Substrate (Bio-Rad Laboratories) was used as a detection system. Pixel intensity quantification was performed with Image Lab software (Bio-Rad Laboratories).

### ELISA receptor phosphorylation assay

Detection of receptors and phospho-receptors was performed in a semi-quantitative way by using a Human Phospho-VEGF R2/KDR DuoSet IC ELISA kit and Human Phospho-PDGFR-β, DuoSet IC ELISA kit (DYC1767-2 and DYC1766-2, R&D Systems) according to the manufacturer’s instructions. For hSDF-1, Phospho-CXCR4 (Ser339) Colorimetric Cell-Based ELISA Kit (OKAG01771, Aviva Systems Biology, San Diego, CA, USA) was used according to the manufacturer’s instructions. HUVECs were used to test the recombinant hVEGF-165 and hSDF-1 bio-functionality, whereas hASCs were used for the hPDGF-B experiments. As positive controls, cells were stimulated with either 50 ng·ml^−1^ recombinant hVEGF-165 for 5 min (Peprotech, Rocky Hill, NJ, USA), 20 ng·ml^−1^ recombinant hPDGF-B for 5 min (Peprotech, Rocky Hill, NJ, USA) and 2 ng·ml^−1^ recombinant hSDF-1 for 10 min (Peprotech, Rocky Hill, NJ, USA). Concentrated protein supernatants of the genetically modified or recipient strains had either the same amount of total protein or the same amount of recombinant protein according to the experiment.

### Angiogenesis tube formation assay

For preparation of conditioned medium for the angiogenesis tube formation assay, recipient (UVM4 and UVM11) and transgenic *C. reinhardtii* cells were cultured (alone or in co-cultivation) in volumes that were adjusted to maintain the same cell density in all experimental samples (e.g. double volume when co-culturing two strains). Supernatants were collected from the cultures at a cell density of 1·10^7^ cells·ml^−1^ and mixed in a 1:1 ratio with AIM-V medium (Life Technologies, NY, USA). Commercially available recombinant growth factors were used as positive controls at the following concentrations: hVEGF-165, 30 ng·ml^−1^; hPDGB-B, 20 ng·ml^−1^ and hSDF-1, 10 ng·ml^−1^. In addition, AIM-V medium was used as a negative control to normalize the results from each independent experiment. Confluent HUVECs were starved (AIM-V medium, Life Technologies) for 24 h prior to experiments. Matrigel (Corning^®^Matrigel^®^ Growth factor reduced, Tewksbury, MA, USA) was thawed on ice overnight, used to coat μ-Slide Angiogenesis (Ibidi, Martinsried, Germany) and allowed to polymerize for 1 h at 37 °C. Then 10-μL aliquots of HUVECs (1·10^6^ cells·ml^−1^) were seeded on the Matrigel-coated slide, 40 μL of conditioned or control medium was added, and the slide was incubated for 4 to 6 h (37 °C, 5 % CO_2_). Finally, viable cells were distinguished from dead cells by staining with calcein-AM/propidium iodide (1.6 μM calcein-AM, 3 μM propidium iodide), which was directly added to each well and incubated for 5 min at 37°C before imaging the cells on a fluorescence microscope (Axiovert 25, Carl Zeiss AG, Oberkochen, Germany). Loop formation was quantitatively determined by computer-assisted image analysis using ImageJ software (Schneider, Rasband, & Eliceiri, 2012).

### Statistical analysis

All results presented were obtained from at least three independent experiments and are expressed as means ± standard deviation. To determine the statistical significance of differences between groups, one-way ANOVA tests were performed using the GraphPad Prism 8.0 software (San Diego, CA, USA). Differences between means were considered significant when p ≤0.05.

## Results

### Generation and molecular characterization of transgenic Chlamydomonas strains expressing hVEGF-165

The use of *C. reinhardtii* for the development of photosynthetic therapies revealed the unique potential of transgenic microalgae as carriers of both oxygen and bioactive molecules (Chávez et al. 2016; Centeno-Cerdas et al. 2018). To establish an optimized strategy for efficient expression of human growth factors in algae, we set out to compare different combinations of *C. reinhardtii* strains optimized for transgene expression as well as two expression vector systems. Using the human hVEGF-165 transgene as a case study, the vectors pBC1 and pOpt, and the recipient strains UVM4 and UVM11 were evaluated to find the most efficient vector-strain combination. To compare the efficiency of both expression vectors, the vector pOpt_VEFG-165 was constructed using the codon-optimized hVEGF cDNA sequence inserted in the previously reported vector pBC1_VEGF-165 (Chávez et al. 2016). In addition, a variant of hVEGF-165 containing a short affinity tag (streptavidin-tag) to facilitate purification was designed and cloned into the pOpt construct, yielding pOpt_VEGF-165-strep (Fig. 1). Furthermore, we compared the levels of the recombinant proteins produced in the host strains UVM4 and UVM11 (Neupert et al. 2009). For each construct, the respective recipient strain is indicated by the suffix “-4” or “-11”. Finally, the previously generated UVM4-derived pBC1_V transgenic strain (Chávez et al. 2016) was used as a reference, giving a total of six different construct-strain combinations (pOpt_Vs-4, pOpt_Vs-11, pOpt_V-4, pOpt_V-11, pBC1_V-4, pBC1_V-11).

No remarkable differences were observed in transformation efficiencies among constructs or strains. Nevertheless, upon screening of about 200 transformants for each strain-construct combination, a high degree of variation in the numbers of transformants that had integrated the complete hVEGF-165 expression cassette was observed (Table 1). These differences were even more pronounced with respect to the fraction of transformants that secreted the recombinant protein into the medium in detectable amounts. In that case, the pBC1 vector in combination with the strain UVM11 (pBC1_V-11) was found to yield the highest proportion of hVEGF-165-secreting transgenic clones (58% of the transformants). Furthermore, in terms of the range of yields of secreted protein (Supp. Fig. S1), only 18.4 % of the transgenic UVM11 clones generated with the pOpt_VEGF-165 construct secreted more than 0.1 fg hVEGF-165 per cell (Fig. S1A), whereas more than 90% of the clones obtained with the pBC1_VEGF-165 construct exceeded this threshold (Fig. S1B). For each construct-strain combination, the clone that secreted the highest amount of growth factor in this initial ELISA-based screen was selected for all further experiments.

**Table 1.** Overview of the *C. reinhardtii* transgenic strains transformed with the indicated plasmids carrying hVEGF expression cassettes. The numbers of colony-forming clones that were obtained upon transformation and subsequent selection on paromomycin (Transformants) are shown. These clones were screened for integration of the hVEGF-165 transgene into the nuclear genome by PCR (Transgenic) and then tested by ELISA for the ability to secrete the recombinant growth factor into the medium (Secreting). The relative number of transgenic and secreting clones in relation to total number of transformants is given in brackets.

|  | pOpt_Vs-4 | pOpt_Vs-11 | pOpt_V-4 | pOpt_V-11 | pBC1_V-11 |
| --- | --- | --- | --- | --- | --- |
| **Transformants** | 220 | 250 | 220 | 213 | 253 |
| **Transgenic** | 112 (51 %) | 210 (84 %) | 147 (67 %) | 114 (54 %) | 171 (68 %) |
| **Secreting** | 35 (16 %) | 99 (40 %) | 39 (18 %) | 28 (13 %) | 147 (58 %) |

For these hVEGF-165 clones, we used Southern blotting to characterize the numbers of transgene copies inserted into the nuclear genome (Supp. Fig. S2). Almost all were found to carry a single integrated copy of the transgene. The only exception observed was one pOpt_Vs-11 clone, which appeared to have integrated two copies of the transgene into the nuclear genome. Next, levels of transgene expression were measured by Northern blot analysis (Fig. 2A). The two pBC1-derived clones showed the highest mRNA accumulation, with pBC1_V-11 expressing significantly higher levels than the pOpt_V-derived clones. Moreover, while expression levels from pOpt_Vs-4, pOpt_V-4 and pOpt_V-11 were comparable, almost no hVEGF mRNA was detectable in the pOpt_Vs-11 strain. A slight difference in size was observed between the pOpt_V- and pBC1_V-derived transcripts, which was probably caused by the use of different regulatory elements for transgene expression in the two vector systems (expected transcript sizes: pOpt_VEGF-165 1.2 knt, pBC1_VEGF-165 1.4 knt). Altogether, the results obtained suggest that in comparison to the pOpt expression cassette the pBC1 system facilitates the generation of higher numbers of transgenic clones with efficient hVEGF-165 expression.

**Figure 2:**
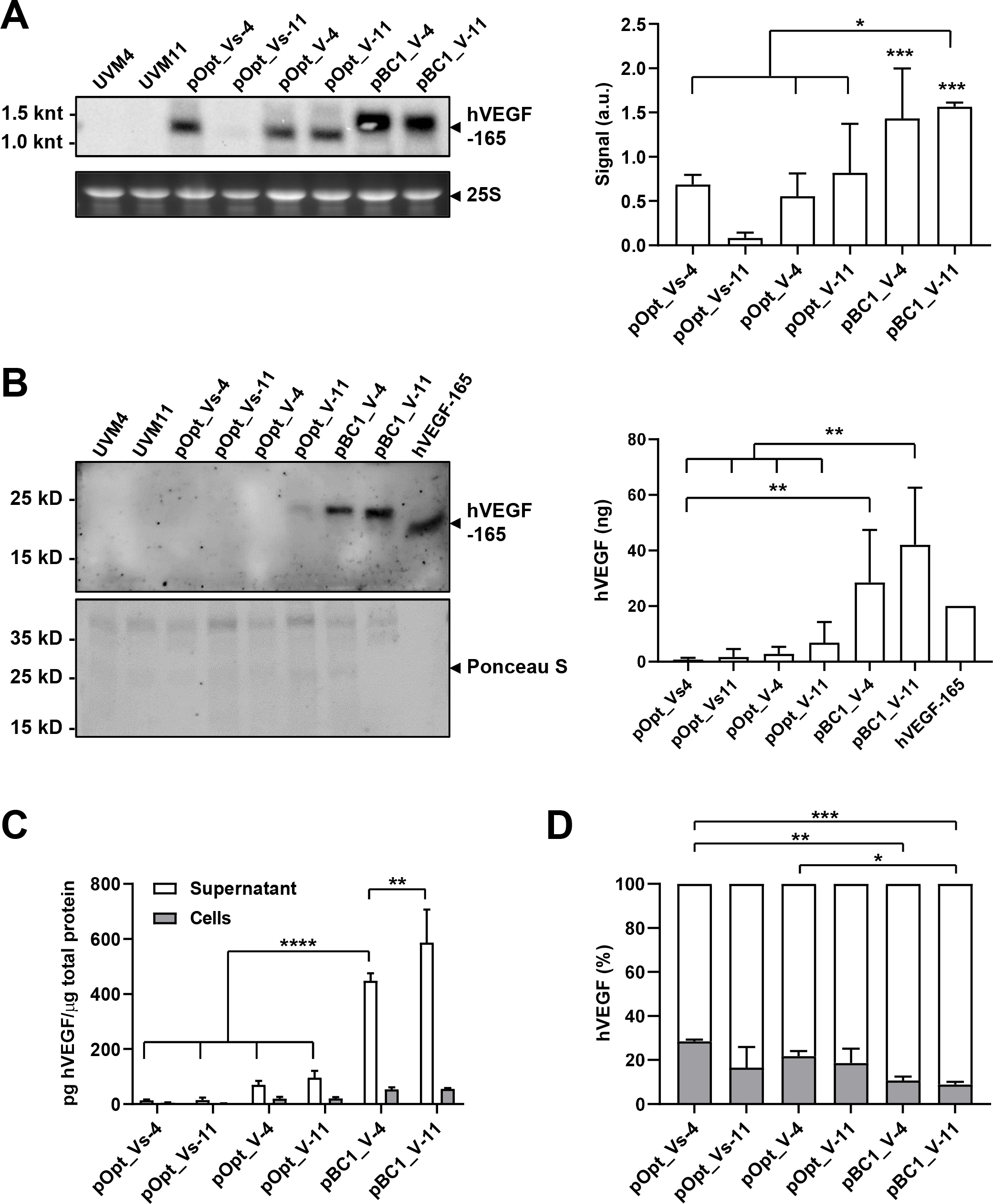
Analysis of hVEGF-165 expression in *C. reinhardtii*. (A) Steady-state levels of the transgene transcript were determined by Northern blot analysis. RNAs extracted from recipient strains (UVM4, UVM11) were used as negative controls to test the specificity of the hVEGF-165 mRNA labelling probe. Chemiluminescence signals were digitally quantified, normalized to a loading control (ethidium bromide-stained 25S ribosomal RNA band after electrophoresis of total RNA, lower panel) and the means of three independent experiments are depicted (right panel). mRNA accumulation was significantly higher in the pBC1-derived clones than in the worst-performing pOpt clone (pOpt_Vs-11). (B) Western blot analysis of hVEGF-165 secreted by each *C. reinhardtii* strain. Volumes of concentrated culture supernatant containing equal amounts of protein (15 μg) were fractionated by SDS-PAGE under reducing conditions. The arrowhead indicates recombinant hVEGF-165, while the lower panel shows Ponceau S staining of the blotted samples (Ponceau S). The amount of hVEGF-165 in the samples (left panel) was determined with reference to a commercially available recombinant hVEGF-165 (20 ng) expressed in *E. coli*. (C) Recombinant hVEGF-165 concentrations in the cell lysates (cells) and algal culture medium (supernatant) were measured by ELISA. Results were normalized to the total protein concentration in the samples. The pBC1-derived strains showed significantly higher yields of secreted protein than any of the others. (D) Secretion yields were calculated from the protein concentrations obtained by ELISA (cells: grey; supernatant: white) and proved to be significantly higher for the pBC1-derived clones compared to the worst-performing clone (pOpt_Vs-4). Only significant differences between groups are specified: **p≤0.005, ***p≤0.001, ****p≤0.0001.

### Impact of transformation vectors and *C. reinhardtii*-strains on recombinant hVEGF-165 yield and bioactivity

In agreement with the observed transcript levels, secreted recombinant protein was only detectable by Western blot analysis in culture supernatants obtained from the pBC1-derived clones, in which a single band of about 20 kD, corresponding to the expected size of the hVEGF-165 monomer, was recognized by the specific anti-VEGF monoclonal antibody (Fig. 2B). The recombinant growth factor expressed in *C. reinhardtii* appeared at a higher molecular weight than the commercially available version of the growth factor expressed in *E. coli*, which might be attributable to post-translational modifications of the protein in *C. reinhardtii* that do not occur in the prokaryotic system. To assess the amount of recombinant protein in the concentrated supernatant, the magnitude of the signal was compared to those given by known concentrations of the commercial hVEGF-165 standard. The calculation led to an estimate of 28.5 ± 18.9 and 42.0 ± 20.5 ng hVEGF-165 for the pBC1-derived UVM4 and UVM11 strains, respectively.

To obtain a more precise quantification of the recombinant protein expressed and secreted by each clone, both culture supernatants and cell lysates were analyzed by ELISA (Fig. 2C). The results show that pBC1_V-11 synthesized more of the protein than any other strain, including the previously characterized pBC1_V-4. Moreover, the levels of secreted growth factor also differed between these two strains, with pBC1_V-11 supernatants showing a small, but statistically significant 1.3-fold increase relative to pBC1_V-4. Also, the secretion ratios calculated from the ELISA data (represented by the fraction of recombinant growth factor in the supernatant, Fig. 2D) showed that, while both pBC1-derived transformants were significantly more efficient than the worst-secreting pOpt_Vs-4 clone, only pBC1_V-11 was statistically better at secreting the non-tagged protein (pOpt_V-4). No significant differences were observed between the UVM4 and UVM11 pOpt-derived clones in either overall yield (Fig. 2C) or secretion efficiency (Fig. 2D). Also, in terms of secretion ratios, no significant differences were observed when one construct was compared with another (pOpt_V-4 vs. pBC1_V-4, pOpt_V-11 vs. pBC1_V11).

We then used both Western blot and ELISA-based semi-quantitative analysis to determine the bioactivity of the recombinant growth factor secreted by each of the transgenic strains by assessing its ability to activate the vascular endothelial growth factor receptor 2 (VEGFR-2). Human endothelial cells were stimulated with concentrated supernatants from each transgenic strain, which either contained the same amount of total protein (Fig. 3A, B) or the same amount of recombinant growth factor (Fig. 3C, D), to induce the dimerization and autophosphorylation of VEGFR-2 upon binding of hVEGF-165. The results showed that the strep-tag version of hVEGF-165 was not able to activate the receptor at all, as receptor phosphorylation upon stimulation did not differ significantly from the negative controls in any of the four assays. The pBC1-coded hVEGF-165 showed the highest receptor activation capacity (Fig. 3C, D). Furthermore, its activity was comparable to that of the commercially available protein in the ELISA-assay when equal amounts of bacterial or algal-derived hVEGF-165 were used, as no significant differences were found among these three experimental groups (Fig. 3D; pBC1_V-4, pBC1_V-11, c+). Interestingly, although statistical analysis failed to reveal any significant differences between the pOpt-encoded and pBC1-encoded hVEGF-165 in the Western blot assay (Fig. 3C), such differences were found when the samples were analyzed by ELISA (Fig. 3D), which suggests that there are differences in biological activity between the products synthesized by the two expression systems. In view of the results obtained for the production of hVEGF-165 from the analyzed algal strains, the use of the pBC1-vector system and the recipient strain UVM11 emerged as promising tools for the production of biologically active hVEGF-165 and potentially other growth factors in *C. reinhardtii*.

**Figure 3:**
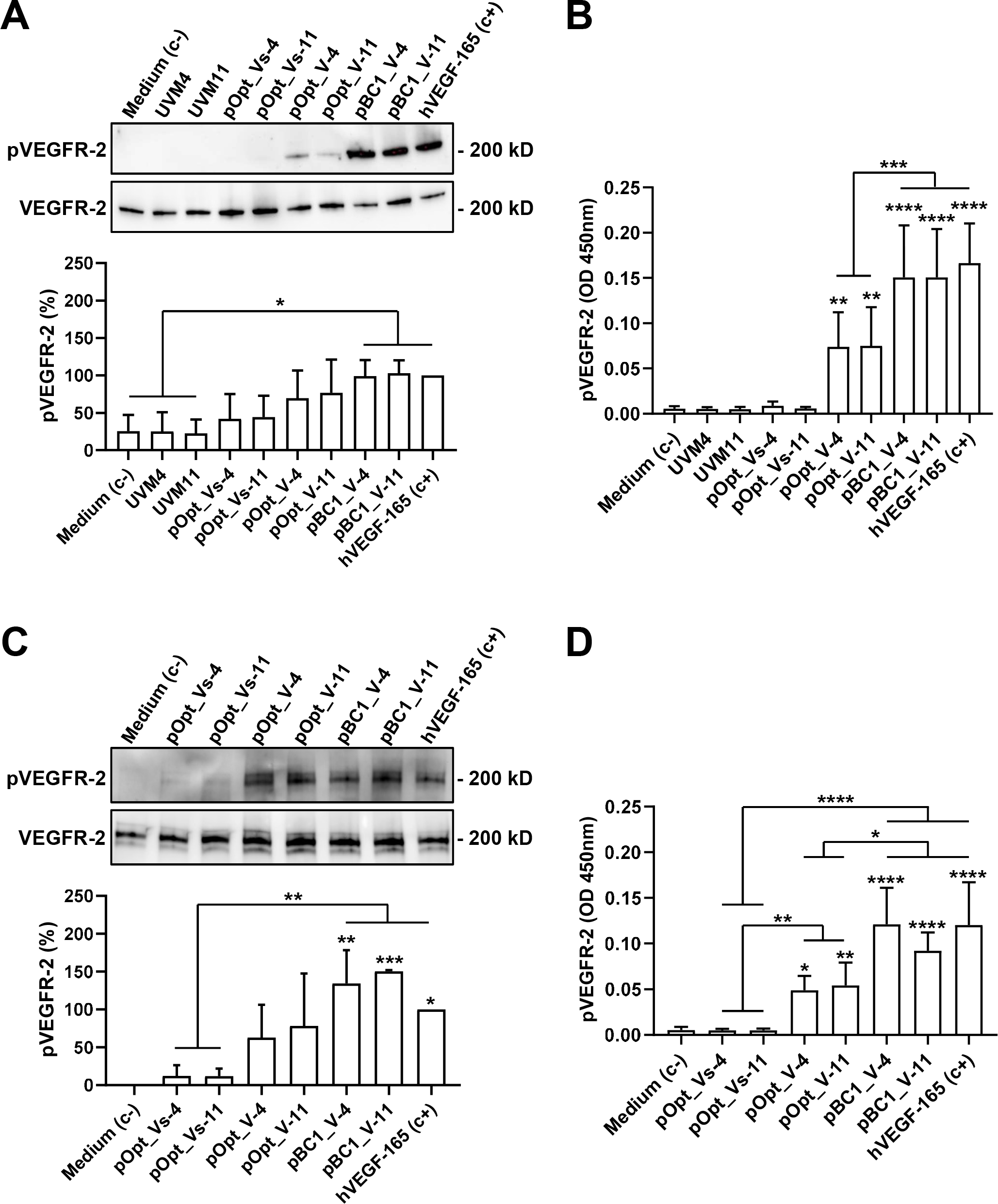
Bioactivity of hVEGF-165 expressed by *C. reinhardtii*. The biological activity of the secreted recombinant growth factors was evaluated for each transgenic strain based on its ability to induce autophosphorylation of VEGFR-2 (pVEGFR-2) as quantified by Western blot analysis (A, C) and ELISA (B, D). For this, HUVECs were stimulated with volumes of concentrated algal culture supernatant that corresponded to either equal amounts of total protein (A, B) or equal amounts of recombinant hVEGF-165 (C, D). The level of receptor phosphorylation (pVEGFR-2 (%)) was calculated based on the signals for the phosphorylated (pVEGFR-2) and total receptor (VEGFR-2) obtained from the Western blot, and the pVEGFR-2-specific semi-quantitative signal measured by absorbance (OD450nm) in the ELISA assay. Statistical analysis was performed with respect to the negative control (starvation AIM-V medium) and to determine differences between strains. A commercially available recombinant growth factor (hVEGF-165) was used as a positive control (c+, 50 ng·ml^−1^). Only significant differences between groups are specified: *p≤0.05, **p≤0.005, ***p≤0.001, ****p≤0.0001.

### Expression of the human pro-angiogenic growth factors hPDGF-B and hSDF-1 using the UVM11 strain and the pBC1 vector system

With the aim of improving the yields of hPDGF-B and hSDF-1 produced by the two UVM4-derived *C. reinhardtii* transgenic strains (pBC1_P-4 and pBC1_ S-4) generated previously (Centeno-Cerdas et al. 2018) and assert the robustness of the identified pBC1-UVM11 system, transgenic UVM11 strains for the expression of these growth factors were also created (pBC1_P-11, pBC1_S-11). Surprisingly, although the rates of successful transgene integration were comparable to that achieved with the pBC1_VEGF-165 construct (see Table 2), the percentage of secreting transgenic clones obtained with pBC1_P-11 and pBC1_S-11 was remarkably low (pBC1_P-11, 16%; pBC1_S-11, 18%; as against pBC1_V-11, 58%; see Tables 1 and 2). Furthermore, in the case of pBC1_P-11 and pBC1_S-11, the proportion of transformants that had integrated the complete expression cassette and were capable of expressing high levels of the recombinant growth factors was very low (75^th^ percentile; pBC1_V-11, 10 of 213; pBC1_P-11, 1 of 218; pBC1_S-11, 7 of 259; see Suppl Figs. S1 and S3).

**Table 2.** Overview of the *C. reinhardtii* transgenic strains transformed with the indicated plasmids carrying hPDGF-B or hSDF-1 expression cassettes. The number of colony-forming clones that were obtained upon transformation and subsequent selection on paromomycin (Transformants) are shown. These clones were screened for the presence of human transgenes in the nuclear genome by PCR (Transgenic), and then tested by ELISA for the secretion of recombinant hPDGF-B or hSDF-1 into the medium (Secreting). The relative number of transgenic and secreting clones in relation to total number of transformants is given in brackets.

|  | pBC1_P-11 | pBC1_S-11 |
| --- | --- | --- |
| **Transformants** | 218 | 259 |
| **Transgenic** | 158 (72 %) | 200 (77 %) |
| **Secreting** | 34 (16 %) | 47 (18 %) |

Nevertheless, a clone was obtained that was able to synthesize and secrete 5.1 times more recombinant hPDGF-B per μg total protein than the previously generated transgenic clones (Fig. 4A), and 8.4 times more when normalized to cell number (Fig. 4B). Furthermore, hPDGF-B secretion into the culture medium was also improved in the UVM11-derived strain, albeit still far below the secretion ratio achieved with the pBC1_V-11 clone (Fig. 2D: pBC1_V-11: 91.2 ± 1.3 %, Fig. 4C: pBC1_P-11: 34.2 ± 3.0 %). Interestingly, there were significant differences in biological activity between the hPDGF-B proteins expressed by the UVM4- and UVM11-derived strains, as shown by their respective capacity to induce autophosphorylation of the platelet-derived growth factor receptor-B (PDGFR-B) (Fig. 4D). Here, mesenchymal stem cells were stimulated with concentrated pBC1_P-4 and pBC1_P-11 culture supernatants, which were diluted to the same growth factor concentration (15 ng·ml^−1^) in human cell culture medium, and compared to the same amount of commercially available hPDGF-B. While all three recombinant growth factors triggered receptor phosphorylation, the results show a higher activity for the protein secreted by pBC1_P-11; however, this increase was minor in comparison to the activity of the hPDGF-B standard used as a positive control.

**Figure 4:**
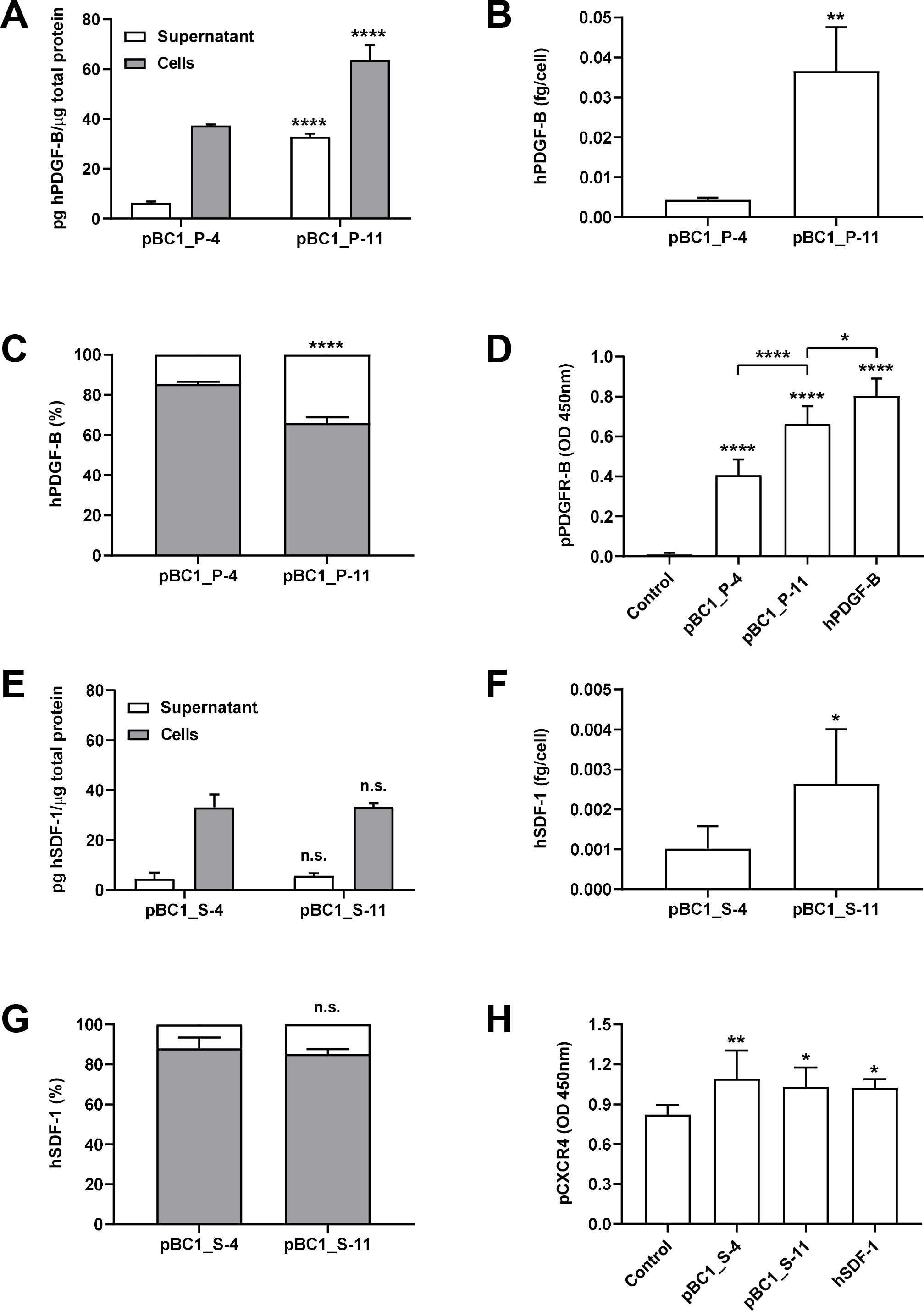
hPDGF-B and hSDF-1 expression in *C. reinhardtii*. Concentrations of hPDGF-B and hSDF-1 synthesized by transgenic *C. reinhardtii* clones were measured by ELISA in both cell lysates and samples of culture medium. Results are normalized to the total protein content (A, E) and cell number (B, F) of the cultured samples. Secretion rates were calculated based on the results obtained by ELISA (C, G). Bioactivity of the secreted hPDGF-B and hSDF-1 was measured through receptor-specific ELISA-based assays, and the phosphorylated forms of the respective receptors (pPDGFR, pCXCR4) were semi-quantitatively determined based on a colorimetric reaction (OD 450nm). For this, human cells (ASCs for hPDGF-B, HUVECs for hSDF-1) were stimulated with equal amounts of the *C. reinhardtii-*secreted or the commercially available recombinant growth factor. * p≤0.05, ** p≤0.01, *** p≤0.001, **** p≤0.0001.

Somewhat surprisingly, in the case of hSDF-1, the best secreting UVM11-derived strain was no better than the previously obtained pBC1_S-4 strain. The results show only statistical differences in the amount of recombinant growth factor secreted per cell, where a 2.6-fold increase with the UVM11 derived strain was estimated. However, even though secretion ratios for both clones were very poor (Fig. 4G; pBC1_S-4: 12.0 ± 5.5 %, pBC1_S-11: 14.8 ± 2.4 %), both recombinant growth factors displayed a comparable bioactivity to the commercially available recombinant hSDF-1, since they were able to induce autophosphorylation of the chemokine receptor type 4 receptor (CXCR4, also known as stromal cell-derived factor 1 receptor) at similar rates (Fig. 4H). All in all, the use of the pBC1-vector and the UVM11 strain appreciably support higher synthesis yields and bioactivity of the recombinant growth factors synthesized in *C. reinhardtii*, though the effect was more pronounced for hPDGF-B than for hSDF-1.

### Combinatorial application of transgenic strains expressing hVEGF-165, hPDGF-B, or hSDF-1 have a potentiating effect on endothelial cell tube formation

As a final assay to demonstrate the applicability of selected transgenic strains, the cumulative effect of the three growth factors on endothelial tube formation and vessel anastomosis was tested in an *in-vitro* angiogenesis assay (Fig. 5). Endothelial cells were seeded on a surface coated with extracellular matrix and stimulated with supernatants obtained from UVM4- and UVM11-derived hVEGF-165 strains, either alone or in combination with the UVM11-derived hPDGF-B and/or hSDF-1 expressing algae. A cumulative potentiating effect was observed when all three recombinant growth factors were present, as shown by the significantly higher numbers of vessel loops that formed compared to the recipient-strain controls (Fig. 5: UVM4, UVM11). This was not achieved by hVEGF-165 alone, regardless of its algal (Fig. 5: pBC1_V-4, pBC1_V-11) or bacterial origin (Fig. 5: Control). Unexpectedly, even though no differences were observed among the pBC1-derived clones in the VEGFR-2 receptor bioactivation assay (Fig. 3), significant differences to the negative controls were observed in this angiogenesis assay only when pBC1_V-4 was combined with the pBC1_P-11 and pBC1_S-11 strains. Interestingly, the combination of hVEGF-165 and hSDF-1 appeared to be more effective than hVEGF-165 and hPDGF, despite the low amounts of recombinant growth factor expressed by pBC1_S-11 strains. These results demonstrate how optimized transgenic *C. reinhardtii* strains may be combined and cultured together to produce stable recombinant growth factor cocktails with high biofunctional activity, in this case, a greater pro-angiogenic effect.

**Figure 5:**
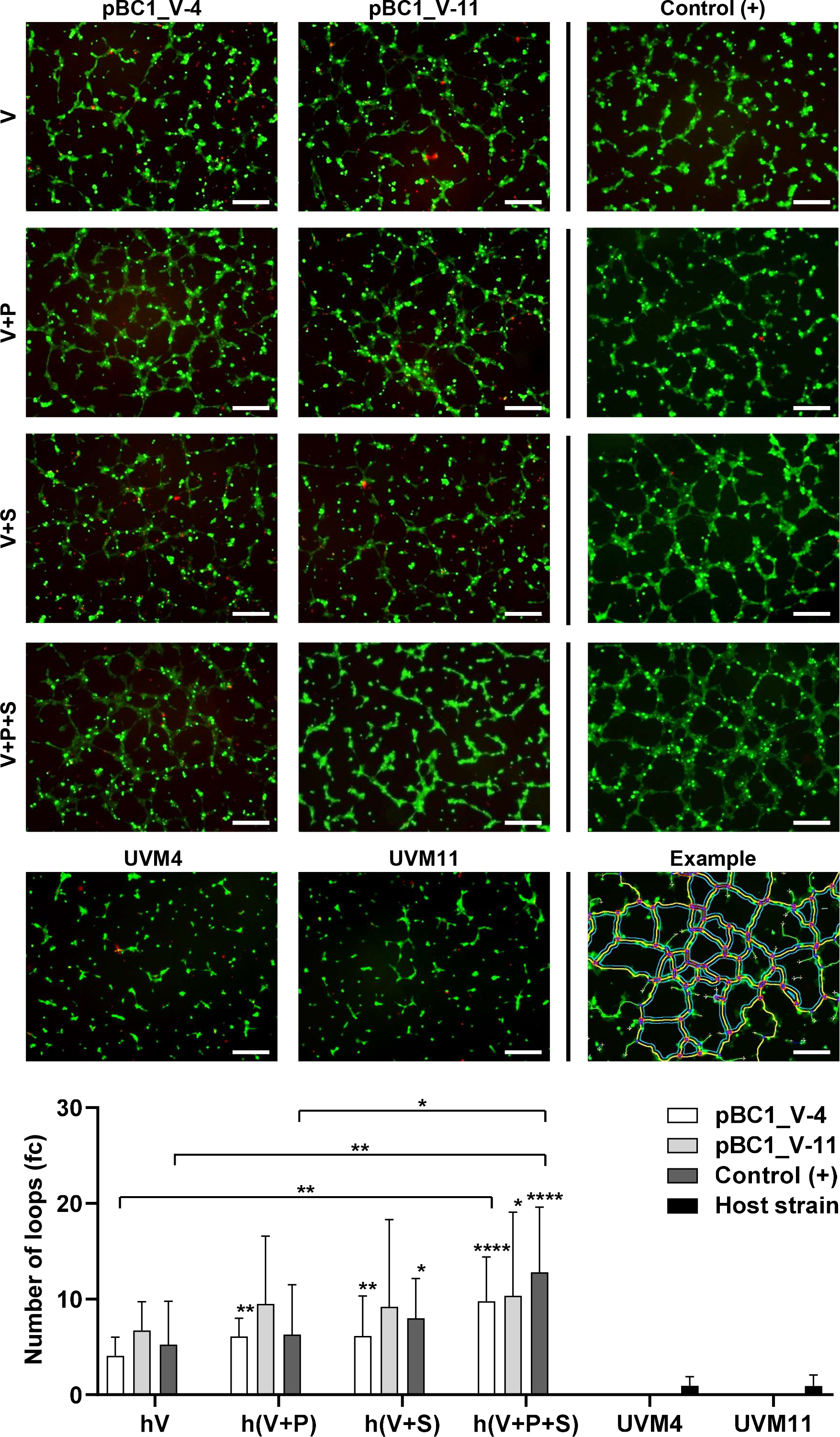
Angiogenesis assay. The pro-angiogenic effect of conditioned media prepared with concentrated supernatants of hVEGF-165-expressing UVM4 and UVM11 *C. reinhardtii* strains (pBC1_V-4, pBC1_V-11), cultured either alone or in combination with the best-secreting hPDGF-B (pBC1_P-11, V+P) or hSDF-1 (pBC1_S-11, V+S) strains, or with both (V+P+S), was evaluated based on their ability to stimulate vessel-loop formation in human endothelial cells. Conditioned media prepared with the recipient strains (UVM4, UVM11) served as negative control, while conditioned media containing specific concentrations of commercial, bacterially derived recombinant growth factors (hVEGF-165: 30 ng·ml^−1^, hPDGF-B: 20 ng·ml^−1^, hSDF-1: 10 ng·ml^−1^) served as positive controls (Control). The results of three independent experiments were normalized to an internal experimental control using the AIM-V culture media alone and are therefore expressed as fold-change (fc). An example of the results obtained from digital image analysis illustrating the loops quantified for the positive control containing all three growth factors is shown at the bottom right of the panel. Statistical significance with regard to the negative controls (recipient strains) is shown, as are the significant differences observed between the experimental groups and expressed above each set of columns). Only significant differences between groups are specified: *p≤0.05, ** p≤0.01, **** p≤0.0001. The scale bar represents 250 μm.

## Discussion

Efficient expression of nuclear transgenes for recombinant proteins in *C. reinhardtii* is a long-sought goal in the biotechnology of algae. *C. reinhardtii* is regarded as one of the most attractive organisms for recombinant protein expression because its genomes have been fully sequenced and well-established protocols for transformation, mutagenesis and mating are available (Rasala and Mayfield 2015). In terms of proliferation speed, scalability, sustainability and safety, *C. reinhardtii* offers many advantages and, aided by its status as a GRAS organism (i.e., generally recognized as safe), it has proved to be a reliable option for the production of almost 50 different recombinant proteins so far (Rasala et al. 2014; Dyo and Purton 2018). In previous work, we attempted to develop strains that could express and secrete human growth factors *in situ* to promote angiogenesis and wound healing in various biotechnological applications (Chávez et al. 2016; Centeno-Cerdas et al. 2018). These studies showed *C. reinhardtii* to be capable of expressing functional recombinant growth factors in both a stable and constant manner. Besides, due to their intrinsic capacity for photosynthesis, the transgenic algae were able to produce oxygen from the biomaterials they were seeded on, hence converting them into the first reported biomedical devices capable of the simultaneous production of oxygen and recombinant biomolecules. However, differences in expression rates between transgenic strains expressing hVEGF-165, hPDGF-B and hSDF-1 alerted us to the need to standardize the expression platform, in order to expand the potential of photosynthetic gene therapy. In the present investigation, we set out to identify the best combination of expression elements and *C. reinhardtii* strains from those described in three previous studies on high-efficiency recombinant protein expression.

We began by testing the pBC1 (Fischer and Rochaix 2001) and pOpt (Lauersen et al. 2013) vector systems. Both vectors allow the insertion of transgene coding sequences into expression cassettes for expression in the nucleus of the host, but they differ in the promoters and regulatory signals they contain. Moreover, two mutant strains UVM4 and UVM11 (Neupert et al. 2009), which have shown a capacity for enhanced transgene expression were compared as hosts, in an attempt to find a strain that consistently produced larger amounts of recombinant growth factors than the strains previously reported by our group. The results revealed that the efficiency of transgene integration into the nuclear genomes of the hosts tested varied between strains (Table 1). Also, compared to the 58% hVEGF-165-secreting pBC1_V-11 clones, only 13 % of the pOpt_V-11 clones selected on paromomycin secreted the recombinant protein, which is considerably less than the 30±5% reported previously (Lauersen et al. 2015). Besides, significant differences in levels of transgene transcription were observed among the strains generated.

The *PsaD* and *HSP70A/RBCS2* promoters are the most effective constitutively active promoters for nuclear transgene expression in *C. reinhardtii* described to date. In the case of *PsaD*, all regulatory sequences required for high level expression have been described to lie within its flanking regions (Fischer and Rochaix 2001), whereas in the first version of the pOpt construct (Lauersen et al., 2013), no further intronic sequences were introduced to effectively produce high amounts of recombinant proteins. Yet, our results show an advantage of using the pBC1-based system to drive the expression of intron-less transgenes. In this regard, it has been suggested that differences between the design of the cDNA introduced and the native exon-intron structure could contribute to poor transgene expression in this organism (Ramos-Martinez et al. 2017; Molino et al. 2018; Baier et al. 2018a). While the proteins encoded by the endogenous genes expressed from the promoters used in our study are very similar in size (PsaD: 21.3 kD; RBCS: 20.6 kD), the exon-intron structure of their genes differs clearly. *RBCS* contains four exons while *PsaD* represents an intron-less gene. Since the hVEGF-165 transgene is derived from a full-length intron-less cDNA, this could explain the higher expression levels of the hVEGF-165 transgene upon integration into the pBC1 vector under control of *PsaD* based regulatory elements as compared to the *HSP70A/RBCS2* based pOpt system.

Interestingly, in the latest version of the modular pOpt vector, all reporter transgenes were interrupted by the second intron of *RBCS2* (Lauersen et al. 2015) leading to an accumulation of secreted reporter proteins of up to ~0.7 mg·l^−1^ under standard cultivation conditions. In this regard, Baier and co-workers recently determined how to design heterologous transgenes with optimally introduced artificial intronic sequences (Baier et al. 2018b). Their results agreed with previous studies suggesting that spliceosome binding motifs in *C. reinhardtii* correlate with the canonical eukaryotic consensus sequence. This finding subsequently led to the development of an algorithm for *in silico* sequence optimization of transgene transcripts that correctly predicts enhanced heterologous gene expression in *C. reinhardtii* (Weiner et al. 2018). These recent strategies could be implemented in the future design of expression cassettes for human transgenes.

In terms of secretion rates, both pBC1-derived strains performed better than any of the pOpt-derived strains (Fig. 2D), although statistically significant differences were only observed when the pOpt_V-4 strain was compared with the pBC1_V-11 strain. This result is unexpected, since the cCA1 secretion signal provided by the pOpt system has been shown to lead to the accumulation of up to 10 mg·L^−1^ of recombinant protein in the supernatant (Lauersen et al. 2013), and it was recently found to be comparable in this respect to the ARS1 and ARS2 secretion signals, of which the latter is used in the pBC1 system (Baier et al. 2018a). On the other hand, similar results to ours have been reported by Molino and co-workers, who found that the use of the cCA1 secretion signal elicited low-level expression of mCherry in *C. reinhardtii* (Molino et al. 2018). The authors suggested that inefficient cleavage of the signal peptide or interferences among the amino acid sequence might impede protein expression. Nevertheless, hVEGF-165 expressed from both pOpt and pBC1 cassettes was able to induce receptor dimerization and phosphorylation, although not with equal efficiencies (Fig. 3D).

A further goal of this work was to develop a strategy for purifying recombinant growth factors secreted by *C. reinhardtii*. Therefore, a strep-tag was fused to the C-terminus of the recombinant hVEGF-165. However, the presence of this sequence negatively affected both transcript and protein accumulation, as well as the bioactivity of the recombinant growth factor (Figs. 2, 3). Similar results were observed in another study (Crozet et al. 2018), where the insertion of 6His- and strep-tags affected the functionality of the fusion protein.

The UVM11-derived strain pBC1_V-11 outperformed the previously reported UVM4-based pBC1_V-4 transgenic strain (Chávez et al. 2016), even though a previous comparison between these two host strains revealed no differences between UVM4 and UVM11 (Kong et al. 2014). Thus, we tested whether this holds also for the UVM4 strains expressing hPDGF-B and hSDF-1 created previously (Centeno-Cerdas et al., 2018). Here, the transformation of UVM11 with the pBC1_PDGF-B and pBC1_SDF-1 constructs led to the recovery of more strains with improved yields of recombinant protein and, in the case of hPDGF-B, significantly increased bioactivity. However, differences were observed in the percentage of secreting clones obtained for each transgene. Thus, the numbers of such clones obtained for hPDGF-B and hSDF-1 were at best only ≈70% as high as for hVEGF-165 (Table 1 and Table 2). Moreover, the numbers of highly expressing clones were ≈10- and ≈2-fold lower than for hVEGF-165, respectively (Supp. Fig. S1 and Supp. Fig. S3). In the study performed by Neupert and co-workers, the authors report a success rate of 100 % for transgene expression (Neupert et al. 2009), which should guarantee that all transformed clones that have integrated the complete transgene cassette can express it. Also, they observed that all transgenic clones had similar levels of recombinant protein, though they warned that mRNA instability and inefficient translation are also important determinants of transgene expression rates. The results obtained here confirm this last observation since protein yields varied significantly for all produced growth factors. On a positive note, a recent study attempted to implement Neupert’s UV-induced mutagenesis strategy to create efficient expression strains from DNA methyltransferase-deficient *C. reinhardtii* mutants (Kurniasih et al. 2016). Here, the authors reported a ratio of almost 74 % highly expressing transformants with one of these mutants, in contrast to the 39 % obtained with UVM4, which suggests the creation of a more robustly expressing strain for transgenes, and another possibility of obtaining strains that produce high levels of hPDGF and hSDF1.

Furthermore, since both hPDGF-B and h-SDF1a seem to be difficult targets for heterologous expression, two recent studies have reported novel strategies to promote a more efficient secretion of microalgae-based recombinant proteins, which might serve to improve the production of extracellular hPDGF-B and hSDF-1. In the first, inspired by a successfully implemented strategy to enhance secretion yields and the stability of recombinant proteins in plant cell cultures, Ramos-Martinez and co-workers showed that linking the N-terminal signal sequence of the metalloprotease gametolysin to the yellow fluorescent protein Venus and fusing a synthetic glycosylated serine-proline tandem sequence to its C-terminus improved its secretion yield by up to 12-fold compared to Venus without the glycomodule (Ramos-Martinez et al., 2017). In the second approach, Baier and co-workers investigated the use of C-terminal fusion peptides to enhance secretion (Baier et al. 2018a). They first identified the secretion signal of an iron assimilatory protein (FEA2) as most effective in targeting the codon-optimized *Gaussia princeps* luciferase (gLuc) into the extracellular space, when compared to the signal peptides of cCA, ARS1 and ARS2. Then, they optimized the *Lolium perenne* ice-binding protein as a fusion protein to increase reporter protein secretion. Finally, they tested the secretion efficiency of the newly created expression cassette with the human epidermal growth factor (hEGF) and reported reaching yields of 0.2−0.25% of the total secreted protein compared to the 0.06 % (Fig. 2C, pBC1_V-11: 586.6 pg/μg protein), 0.003 % (Fig. 4A, pBC1_P-11: 32.9 pg/μg protein) and 0.0006 % (Fig. 4E, pBC1_S-11: 5.8 pg/μg protein) obtained here for hVEGF-165, hPDGF-B and hSDF-1, respectively.

Strikingly, despite the relatively low yields of secreted hSDF-1 and hPDGF-B (relative to hVEGF) obtained in our experiments, the supernatants of co-cultivated strains expressing all three growth factors exhibited significantly higher levels of functionality in angiogenesis assays than did the strain expressing hVEGF alone (Fig. 5). This could have a significant impact as a potential off-the-shelf biomedical application, since it would be possible to combine transgenic *C. reinhardtii* strains to produce defined growth-factor cocktails capable of stimulating specific cellular processes, or even formulations specifically adapted to the needs of the individual patient, once a complete set of transgenic alga strains for every therapeutic peptide on the market is established. Taken together, in this work, we describe an attempt to study some of the key factors required for the efficient secretion of human growth factors by transgenic *C. reinhardtii* strains and discuss elements that could guarantee high recombinant protein expression yields for all the transgenes studied. Our results show the superiority of the pBC1 system for the generation of high-secreting clones, and the greater potential of the UVM11 strain over UVM4 as a host for the secretion of higher yields of human growth factors. However, further efforts to find optimization strategies are required before a standardized protocol capable of generating highly efficient recombinant strains for photosynthetic therapies can be formulated.

## Compliance with ethical standards

### Funding

This work was supported by a grant from the World Bank to MJC (Grant No. 05-15-D) and the Deutsche Forschungsgemeinschaft to JN (SFB TRR175-A06).

### Conflict of interest

The authors declare that they have no conflict of interest.

### Ethical approval

This article does not contain any studies with human participants or animals performed by any of the authors.

### Author contributions

Conceptualization: JN, JTE, MJC; Methodology: MJC, MNC, AVB; Formal analysis and investigation: MJC; Writing - original draft preparation: MJC, MNC; Writing - review and editing: MNC, AVB, JN; Funding acquisition: MJC, JN, TE; Resources: JN; Supervision: JN. All authors read and approved the final manuscript.

## Notes

#### Summary of Updates

Revised version upon the peer-review process.

