## Supplemental information for "Towards a biotechnological platform for the production of human pro-angiogenic growth factors in the green alga *Chlamydomonas reinhardtii*"

**Affiliations:**
[a] Molekulare Pflanzenwissenschaften, Biozentrum Ludwig-Maximilians-Universität München, Planegg-Martinsried, Germany
[b] Biotechnology Research Center, Instituto Tecnológico de Costa Rica, Cartago, Costa Rica
[c] Department of Plastic Surgery and Hand Surgery, University Hospital rechts der Isar, Technische Universität München, Germany
[d] Institute for Biological and Medical Engineering, Schools of Engineering, Biological Sciences and Medicine, Pontificia Universidad Católica de Chile

***Corresponding author**
Prof. Dr. Jörg Nickelsen
Biozentrum LMU München
Grosshaderner Strasse 2-4, D-82152 Planegg-Martinsried
Telephone: +49 (0)89 / 2180-74773

### Supplementary Figure S1:

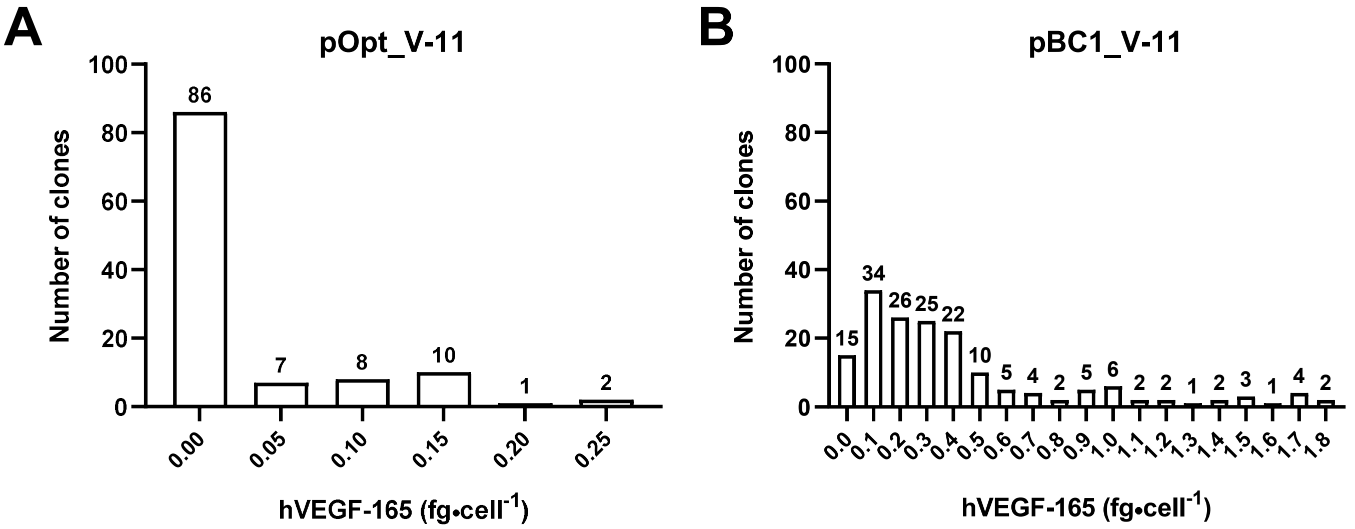


Supp. Fig. S1: Histograms showing the amounts of hVEGF-165 secreted by the transgenic clones generated with the pBC1_V and pOpt_V vectors. Culture supernatants from the UVM11 transgenic clones generated using the pOpt (A) and pBC1 vector (B) were analyzed by ELISA to quantify the concentration of the secreted recombinant growth factor. Data were then ordered to represent the distribution of the clones according to the amount of secreted hVEGF-165.

### Supplementary Figure S2:

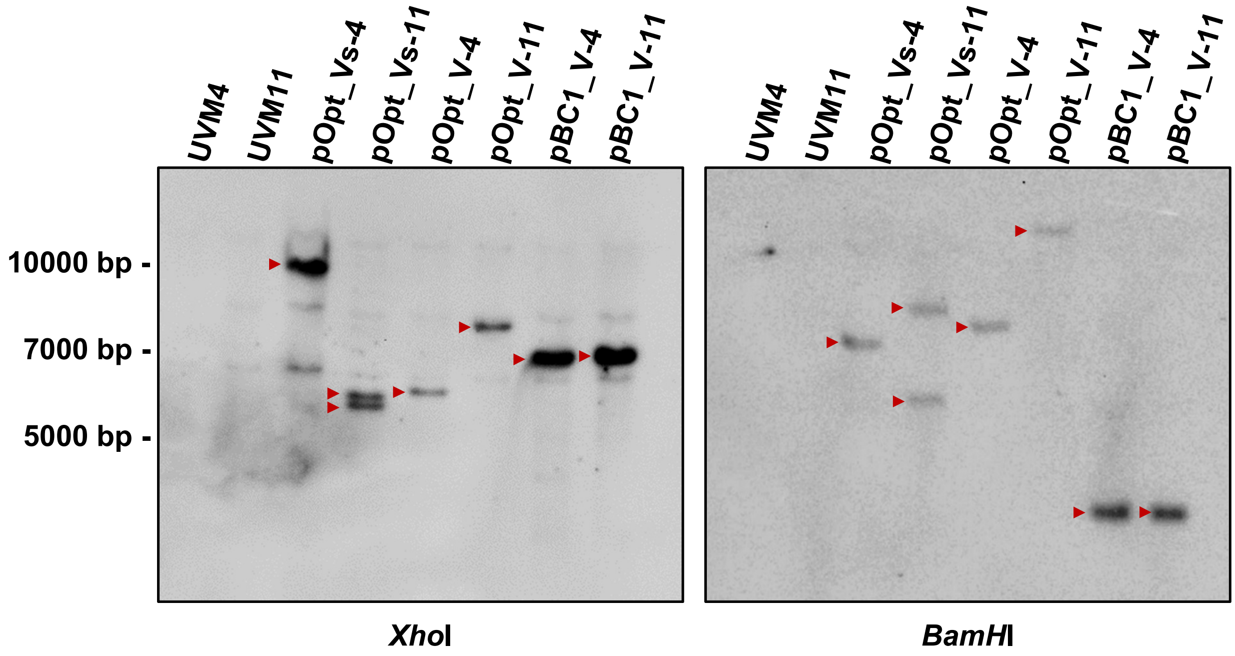


Supp. Fig. S2: Verification of hVEGF-165 transgene integration in the nuclear genome of *C. reinhardtii*. The number of integrated copies of the hVEGF-165 coding transgene was verified by Southern blot analysis. Total DNA was digested with the restriction enzymes *Xho*I and *BamH*I, and size-separated by agarose gel electrophoresis. Recipient strains (UVM4, UVM11) were included as negative controls to test the specificity of the hVEGF-165 labelling probe. Most of the transformants have a single copy of the VEGF transgene. A representative image of one out of three independent experiments is shown.

#### Supplementary Figure S3:

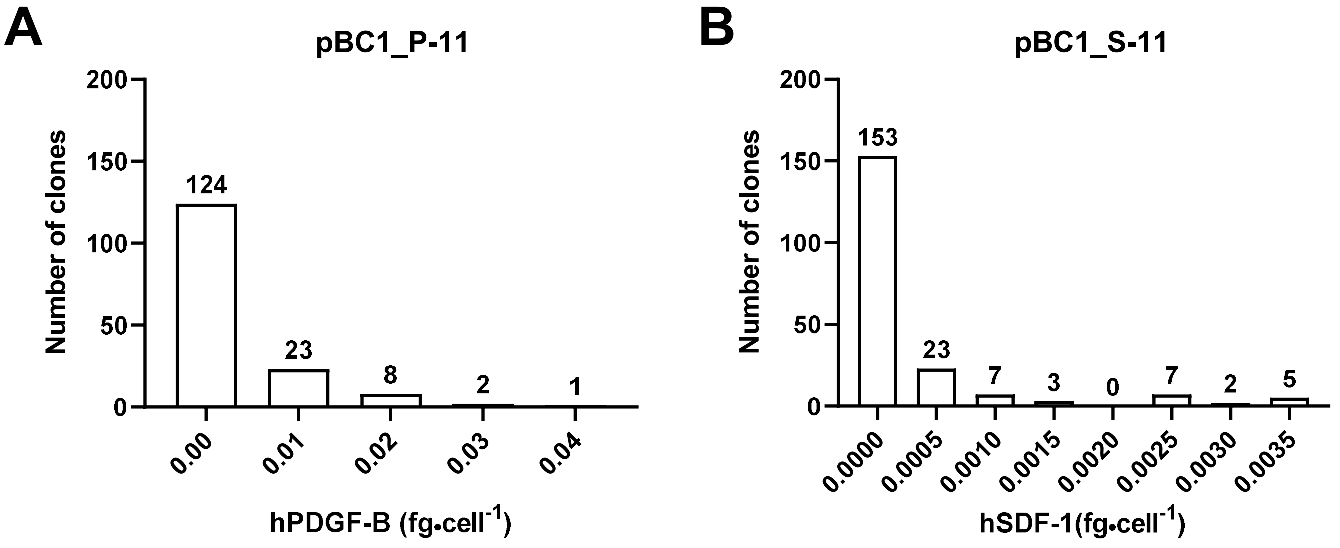


Supp. Fig. S3: Histograms showing the amounts of hPDGF-B and hSDF-1 secreted by the transgenic clones generated with the pBC1_P and pBC1_S vectors. Culture supernatants from the UVM11 transgenic expressing hPDGF-B (A) or hSDF-1 (B) were analyzed by ELISA to quantify the concentration of the recombinant growth factors. Data were then ordered to represent the distribution of the clones according to the amount of secreted growth factors.
